# Rapid 40 kb genome construction from 52 parts

**DOI:** 10.1101/2020.12.22.424019

**Authors:** John M. Pryor, Vladimir Potapov, Nilisha Pokhrel, Gregory J. S. Lohman

**Affiliations:** Research Department, New England Biolabs, Ipswich, MA, 01938, USA

## Abstract

Large DNA constructs (>10 kb), including small genomes and artificial chromosomes, are invaluable tools for genetic engineering and vaccine development. However, the manufacture of these constructs is laborious. To address this problem, we applied new design insights and modified protocols to Golden Gate assembly. While this methodology is routinely used to assemble 5-10 DNA parts in one-step, we found that optimized assembly permitted >50 DNA fragments to be faithfully assembled in a single reaction. We applied these insights to genome construction, carrying out rapid assembly of the 40 kb T7 bacteriophage genome from 52 parts and recovering infectious phage particles after cellular transformation. The new Golden Gate assembly protocols and design principles described here can be applied to rapidly engineer a wide variety of large and complex assembly targets.

## INTRODUCTION

Large DNA constructs (~10-50 kb) containing multiple coding regions are widely used in biotechnology for the development of vaccines, therapeutics, and genetically engineered organisms. In most cases these constructs are too long to be reliably generated by continuous chemical DNA synthesis methodologies or PCR, and thus are typically assembled from constituent DNA fragments over multiple rounds of assembly [1, 2]. Several commonly used methodologies to manufacture large DNA constructs rely on *in vivo* recombination, and these high capacity systems allow researchers to join up to 40-50 DNA fragments in a single assembly round [3, 4]. However, these techniques typically require lengthy protocols with specialized reagents, and are not amenable to the high-throughput workflows relied upon to engineer desired constructs and gene circuits of a smaller size. *In vitro* DNA assembly methodologies are not encumbered by the same limitations; however, these methods typically limit researchers to 5-10 DNA fragments per assembly round. Thus, the use of hierarchical assembly schemes that involve multiple rounds of molecular cloning, construct purification, and sequence verification of donor constructs are typical for assembly of large constructs using *in vitro* assembly methodologies [5–7]. However, recent work by our lab and others suggests that *in vitro* DNA assembly methods may be able to accommodate many additional fragments per assembly round and thus could be used to rapidly engineer large and complex DNA target sequences [8–10].

Golden Gate assembly (GGA), sometimes referred to as Type IIS assembly, is an *in vitro* DNA assembly methodology that utilizes a Type IIS restriction endonuclease to generate DNA fragments with short single-stranded overhangs and a DNA ligase to join the fragments together [11, 12]. This molecular cloning methodology is extremely versatile. Type IIS restriction enzymes cleave outside of their recognition sequence; this allows users to choose the connecting sequences between assembly fragments and has given rise to more than a dozen ‘modular’ cloning systems with prefabricated parts [6, 13–40]. Additionally, the method can also be used to generate DNA targets without the need to introduce unwanted sequence at fusion sites. A significant drawback of GGA is that Type IIS recognition sequences for the enzyme used in the assembly cannot be present in the desired target sequence; however, in most cases this limitation is easily overcome by choosing a Type IIS restriction enzyme with a minimal number of recognition sequences in the desired target and/or introducing silent mutations [41, 42]. Importantly, this method relies on accurate ligation of the single-stranded DNA sequences at fragment fusion sites, as mis-ligation results in improperly ordered fragments and low assembly yield. To avoid erroneous assembly products caused by ligation errors, GGA has typically been limited to ~10 fragments per reaction; however, a few recent studies have reported >20 fragment assembly of constructs up to 10 kb [8, 10, 28]. Recent work by our lab showed that optimization of GGA by data-driven selection of overhang sequences that adjoin assembly fragments, termed Data-optimized Assembly Design (DAD), can significantly expand the limits of this assembly methodology [10].

We have previously shown that using DAD allows for assembly of up to 35 fragments with high fidelity under traditional GGA reaction conditions [10]. Here, to estimate the maximum number of DNA fragments that can be joined in a single GGA reaction, we analyzed the frequency of erroneous fragment assembly under different reaction conditions using a multiplex DNA sequencing assay developed in-house. Our data suggest that >50 fragments can be assembled in one reaction with high fidelity using an optimized reaction protocol or with stringent screening conditions, if optimal junction selection rules are followed (Supplementary Text, Figures S1 and S2, Tables S1 and S2). In the current study we confirmed these predictions in two practical applications: assembly of a 4.9 kb *lac* operon cassette into a destination vector from 52 constituent fragments, and through assembly of the 40 kb T7 phage genome from 52 parts in one reaction and <1 day.

## RESULTS/DISCUSSION

Our data suggest that prolonged static incubation of GGA reactions at 37°C would allow assembly of >50 parts with comparable fidelity to our previously reported 35 fragment assembly produced under traditional 37°C/16°C thermocycling conditions (Supplementary Text, Figures S1 and S2, Tables S1 and S2). To test this prediction we sought to carry out the most complex assembly reaction to-date and clone a 4.9 kb cassette of the *lac* operon into an *E. coli* destination vector from 52 constituent parts in a single assembly round (Tables S3-S4). Importantly, the *lac* operon cassette system used here mimics a traditional cloning reaction wherein, upon transformation of the assembly reaction into *E. coli* cells, we can observe colonies harboring correctly or incorrectly assembled constructs. This test system was engineered to provide a colorimetric readout to differentiate transformants harboring correctly and incorrectly assembled products [8].

DNA fragments comprising the *lac* operon cassette were generated by PCR and assembled in a reaction containing BsaI-HFv2 and T4 DNA ligase at 37°C for 48 hours. Assembly reactions were then transformed into chemically competent *E. coli* cells, and the resulting transformants were scored as having correctly or incorrectly assembled insert sequences. We found that 49% of the observed transformants harbored correctly assembled constructs, slightly higher than our predicted fidelity of 27% (Figure 1, Table S3). The reaction generated >500 transformants with correctly assembled constructs per 100 μL of *E. coli* outgrowth plated, a surprisingly high yield given the complexity of the assembly reaction. To confirm successful assembly of all 52 inserts, constructs were purified from a subset of colonies and analyzed by PCR and Sanger sequencing. All constructs from colonies scored as having correct assemblies were found to have inserts of the anticipated size and sequence, and constructs from colonies scored as having incorrect sequences contained truncated inserts (Figure S3). Taken together, these data show that >50 fragments can be assembled in one reaction using a high temperature reaction protocol and rational junction selection by DAD.

**Figure 1.**
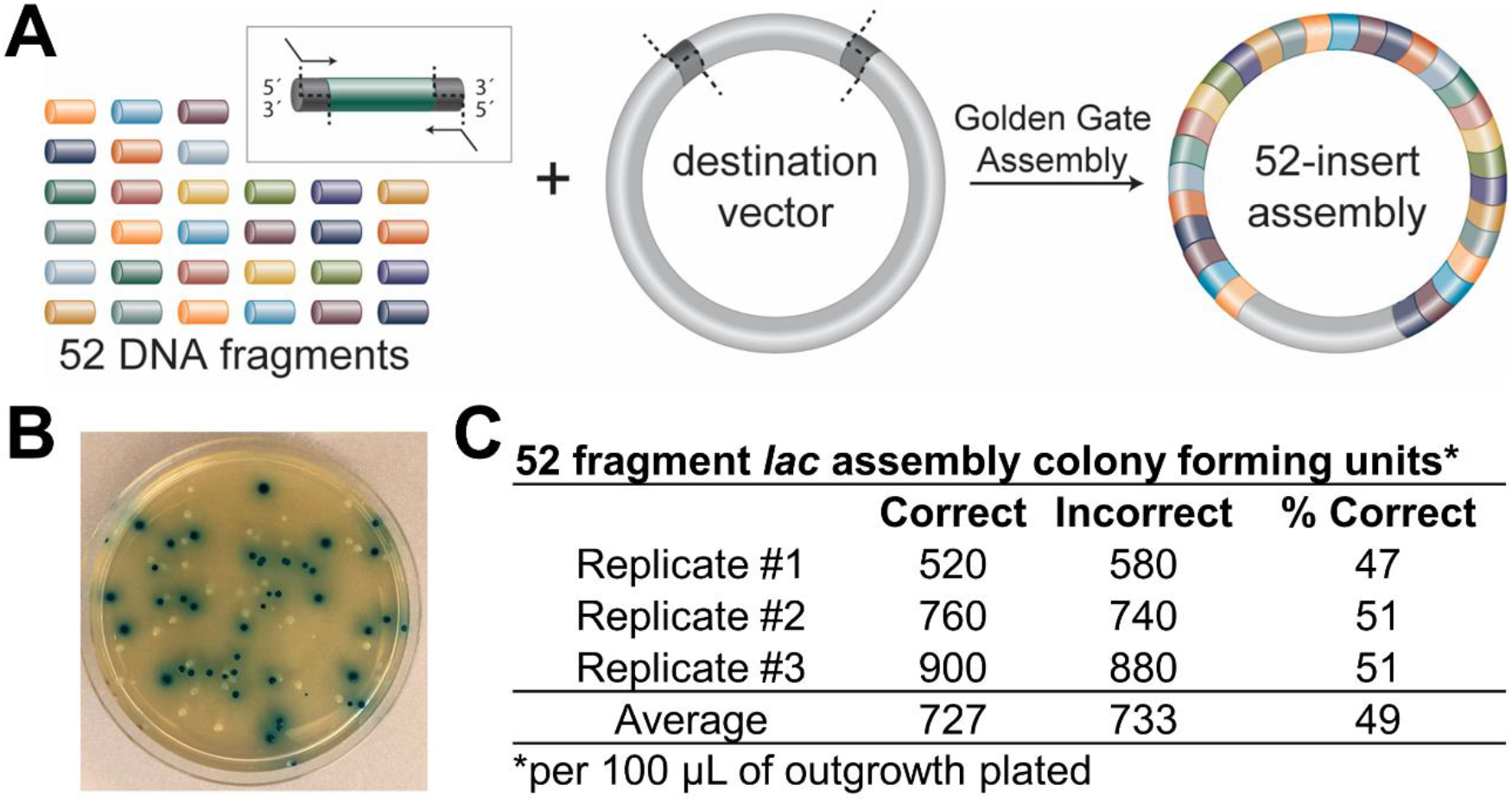
One-pot Golden Gate assembly of 52 fragments into a destination vector. (A) Schematic of the 52 fragment lac operon cassette assembly. Assembly inserts were generated by PCR amplification and assembled into a destination vector containing an antibiotic resistance marker. (B) Example outgrowth plate used for colorimetric scoring by reverse blue-white screening. Correctly assembled 52 insert constructs form blue colonies upon cellular transformation and incorrectly assembled constructs produce white colonies. (C) Results of the assembly reactions. Three replicate experiments were carried out to quantify the number of colony forming units harboring correct and incorrect assembly products per 100 μL of *E. coli* outgrowth plated (0.2 μL of the assembly reaction). On average, 49% of the observed transformants harbored correctly assembled constructs.

Next, we sought to test if using traditional GGA thermocycling protocols would allow for rapid assembly of 50+ fragments, given the use of a more stringent selection system to identify correctly assembled constructs. To test this, we designed a GGA reaction to construct the 40 kb T7 bacteriophage genome from 52 parts (Table S5). Unlike the *lac* cassette system described above, where circularized constructs with deletions produce colonies due to the presence of the vector antibiotic resistance gene, we reasoned that improperly assembled variants of the T7 phage genome would contain significant (100s or 1000s of nt) deletions, and therefore be unlikely to produce viable phage upon cellular transformation. Thus, the T7 genome results in a more stringent selection where most incorrect assemblies will not form plaques. We reconstructed the phage gDNA from PCR amplicons (Table S4). This strategy enabled us to easily introduce 16 silent mutations to remove pre-existing BsmBI Type IIS restriction sites within the phage genome through PCR primers by choosing break points near all native BsmBI sites. These changes serve dual purpose to both permit Type IIS assembly and to act as marker mutations for assembly verification.

The more stringent selection permitted a shorter, cycled assembly protocol that sacrifices fidelity in favor of efficiency and yield. Assembly reactions were carried overnight using a 42°C/16°C oscillating thermocycling protocol with BsmBI-v2 and T4 DNA ligase. The assembly reactions were then transformed into NEB 10-beta electrocompetent cells; successful assembly of the T7 phage genome was assessed by plaque forming assay. Upon transformation we typically observed ~20 bacteriophage plaques/μL of assembly reaction, indicating successful assembly of the phage genome (Figure 2). Several phage plaques were selected for additional screening by plaque PCR and restriction enzyme digest to ensure they contained a complete and correctly ordered copy of the T7 phage genome; all plaques subjected to additional screening contained the expected genome arrangement and harbored the intended silent mutations (Figure S4). Moreover, to ensure the observed phage plaques were the result of *in vitro* assembly and not assembly of the DNA fragments within the *E. coli* by cellular DNA repair mechanisms, we carried out control reactions lacking T4 DNA ligase and did not observe phage plaques upon transformation of these control reactions. Taken together, these results demonstrate that using a high stringency screen allows rapid assembly of >50 DNA fragments under traditional GGA thermocycling conditions with DAD.

**Figure 2:**
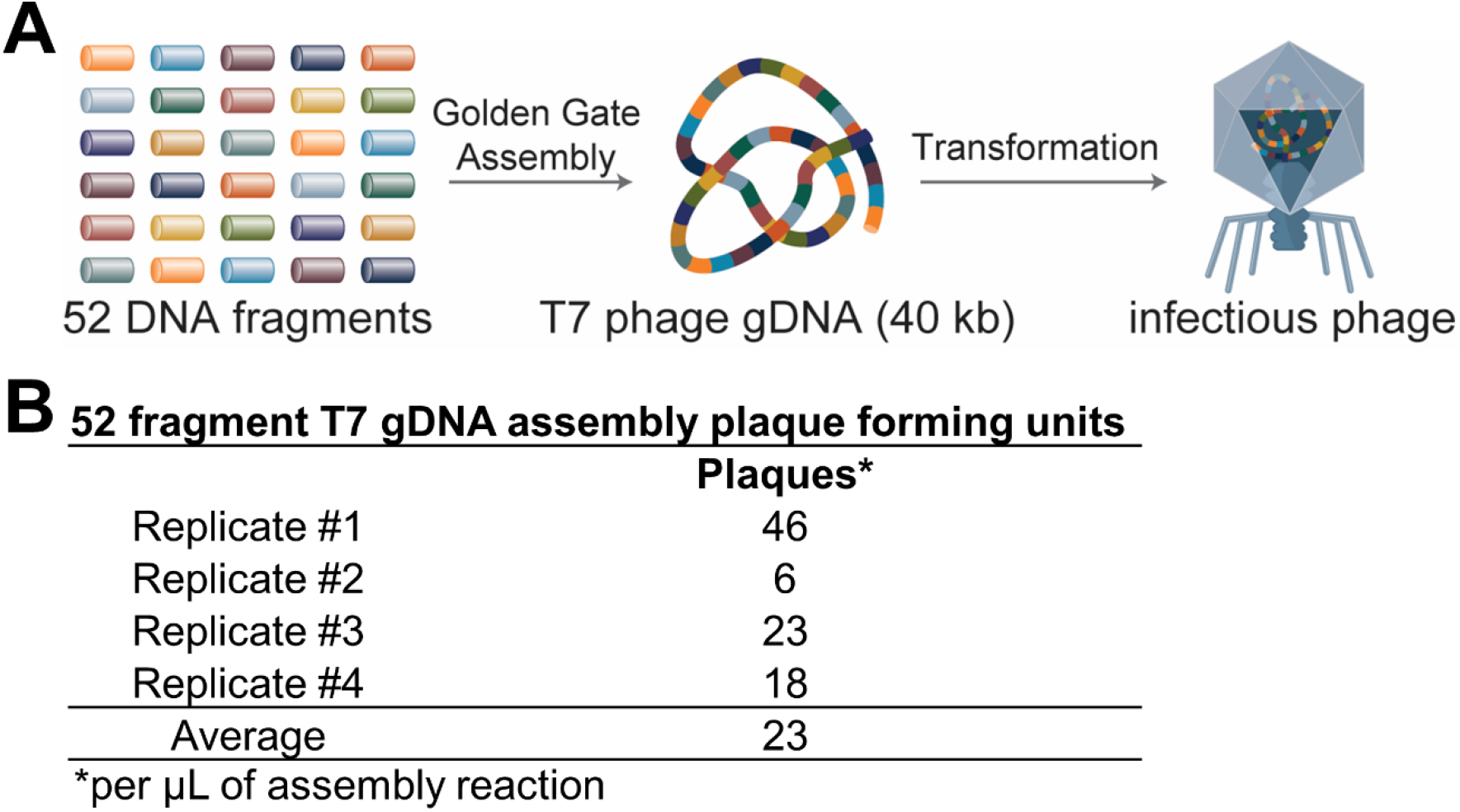
T7 phage genome assembly and infectious phage reconstitution. (A) Schematic of the 52 fragment T7 phage reconstitution experiment. (B) Results of the assembly reactions. Four replicate experiments were carried out to quantify the number of plaque forming units upon transformation of the phage gDNA assembly reactions. On average, 23 plaques were observed per μL of the assembly reaction transformed into *E. coli* cells.

Our results show that >50 fragments can be reliably assembled in a single reaction using GGA with DAD. This finding reduces the number of hierarchical assembly rounds required to produce large constructs by *in vitro* assembly and can be utilized to rapidly assemble entire metabolic pathways and small genomes in a single reaction. Moreover, the large fragment capacity demonstrated here allows for assembly of toxic and/or large DNA constructs from small constituent parts that are easily manipulated and propagated using standard molecular biology techniques, ease of removal of internal Type IIS sites along with fragment generation, and design of large constructs with every ORF on its own fragment. Importantly, this work could facilitate rapid engineering of large DNA targets, as GGA is amenable to automated workflows and supports standardized modular assembly of construct variants with minimal ad-hoc design [43]. In summary, our work demonstrates an efficient and cost-effective means to create and engineer variants of large/complex DNA constructs that are difficult to obtain and manipulate by current cloning and gene synthesis methodologies.

## SUPPLMENTARY TEXT

### The impact of reaction temperature on Golden Gate assembly fidelity

Golden Gate assembly typically utilizes two step cycling protocols, alternating between a 16°C incubation step to maximize DNA ligation efficiency and a 37-42°C incubation step to maximize fragment digestion efficiency. However, we reasoned that omitting the 16°C incubation may increase Golden Gate assembly fidelity, as higher reaction temperatures have been shown to improve DNA ligase fidelity [44]. To test this, we carried out a multiplex high throughput DNA sequencing assay developed in-house to quantify the frequency of Golden Gate assembly errors at 37°C or 42°C, and compared the results to reactions using traditional thermocycling protocols of 37/16°C or 42/16°C (Figure S1). Of note, the reactions carried out at constant incubation temperatures were incubated for an extended duration of 16 hours to compensate for decreased ligation efficiency. We found that the frequency of ligation errors was reduced >2-fold in our sequencing experiments when we omitted the 16°C incubation step, with every mismatch pair appearing less frequently. To asses if more fragments could be assembled in one-step by omitting the 16°C incubation step during Golden Gate assembly, we next estimated assembly fidelity as a function of fragment number as described previously (Figure S2) [10]. We found that the estimated assembly fidelity for traditional 37/16°C or 42/16°C cycling conditions dropped below 10% at 50 fragments, but the 37°C or 42°C static incubation protocols could theoretically allow >50 fragment to be assembled with >40% accuracy. It should be noted that use of the 37°C or 42°C static temperature protocols is likely to require extended duration reactions to compensate for the ligation efficiency loss caused by omitting the 16°C incubation step. Nevertheless, these data suggest that the capacity of Golden Gate assembly reactions could exceed 50 fragments per reaction.

## METHODS

### Reagents and Oligonucleotides

Enzymes, buffers, and media were obtained from New England Biolabs (NEB) unless otherwise noted. Synthetic oligonucleotides were obtained from either Integrated DNA Technologies (IDT) or Sigma Aldrich (Sigma).

### Multiplex DNA Sequencing Assay

Substrates for the DNA sequencing assay were prepared as previously described [8, 10]. Golden Gate assembly reactions (20 μL final volume) to generate sequencing libraries were carried out in 1X T4 DNA ligase buffer by combining: 100 nM of DNA substrate (final concentration) with 2 μL of NEB Golden Gate Enzyme Mix (BsaI-HFv2 or BsmBI-v2). Reactions were carried out for 16 h at 37°C (BsaI-HFv2) or 42°C (BsmBI-v2). Reactions were then quenched by the addition of 25 mM EDTA and column purified (Monarch PCR & DNA Cleanup Kit). The resulting assembly products were further purified to remove un-ligated substrate by treatment with Exonuclease III (50U) and Exonuclease VII (5 U) in 1X Standard Taq Polymerase buffer (final concentration) for 1 h at 37°C in a 50 μL reaction volume. The assembly products were then re-purified using the Monarch PCR & DNA Cleanup Kit and quantified by Agilent Bioanalyzer (DNA 1000).

Pacific Biosciences Single-Molecule Real-Time (SMRT) sequencing was performed as described previously [8, 45]. The libraries were prepared for sequencing using the PacBio Binding Calculator Version 2.3.1.1 and the DNA/Polymerase Binding Kit P6 v2 with a custom library concentration on the plate of 0.3375 nM. Sequencing was carried out using the PacBio RSII instrument with 2 SMRT cells per library and a 3 h data collection time per cell with ‘stage start’ off. Consensus sequences for each assembly product were generated as described previously [8, 45]. Full results from each experiment are supplied in the supporting data files (Tables S1-S2).

### Golden Gate Assembly Reactions

Assembly junction sequences were selected using the optimization described in our previous work [10]. Importantly, these algorithms have been developed into a suite of webtools that can be accessed here: https://goldengate.neb.com. Assembly fragments were generated by PCR (Q5 hot start high-fidelity 2X master mix) with oligonucleotide primers (IDT) and purified using the Monarch PCR & DNA Cleanup Kit. Fragment quality was evaluated using the Agilent Bioanalyzer 2100 and each assembly part was quantified using the Qubit assay (ThermoFisher). Golden Gate assembly reactions (5 μL final volume) were carried out with 3 nM of each DNA fragment and 0.5 μL of the appropriate NEB Golden Gate Assembly Mix in 1X T4 DNA ligase buffer; the BsaI-HFv2 mix was used to reconstitute the *lac* operon cassette and the BsmBI-v2 mix was used to assemble the T7 phage genome. Reactions to reconstitute the *lac* operon cassette were incubated for 48 h at 37°C and then subjected to a final heat-soak step at 60°C for 5 minutes before being incubated at 4°C prior to transformation. Reactions to produce the T7 bacteriophage genome were cycled between 42°C and 16°C for 5 minutes at each temperature for 96 cycles, and then subjected to a 60°C incubation for 5 minutes and finally a 4°C hold until transformation.

### Clonogenic assays

Assembly reactions to reconstruct the *lac* operon cassette were transformed into chemically competent *E*. *coli* cells, and colony forming units were scored as harboring correctly or erroneously assembled constructs by a reverse blue-white screen as described previously [8, 10]. Briefly, transformations were performed using 2 μL of each assembly reaction added to 50 μL of T7 express competent cells as per manufacturer’s instructions. The resulting outgrowth was plated onto agar plates (Luria–Bertani broth supplemented with 1 mg/mL dextrose, 1 mg/mL MgCl2, 30 μg/mL Chloramphenicol, 200 μM IPTG and 80 μg/mL X-gal). Importantly, transformants harboring correctly assembled constructs turn blue after incubation on media containing IPTG and X-Gal, while transformants harboring constructs with assembly errors form white colonies.

### Plaque assays

Reactions to construct the T7 phage genome were transformed into NEB 10-beta cells as per the manufacturer’s instructions, using 1 μL of the reaction mixture into 25 μL of competent cells. The transfection mixture was recovered in 975 μL of NEB 10-beta/stable outgrowth media and then combined with 3 mL of 50°C molten top-agar (Luria broth containing 0.7% agar). Finally, the mixture was plated on LB agar plates and the molten agar was allowed to cool and solidify on the benchtop for 20 m. The resulting pates were inverted and incubated at 37°C for ~5 h until the *E. coli* lawn and phage plaques were visible by eye.

## ACKNOWLEDGEMENTS

We thank Tasha José (New England Biolabs) for providing illustrations as well as and Rebecca Kucera and Eric Cantor (New England Biolabs) for providing reagents. We also thank Katharina Bilotti, Na Ke, Rebecca Kucera, and Eric Cantor (New England Biolabs) for careful reading of the manuscript.

---

**Figure S1.**
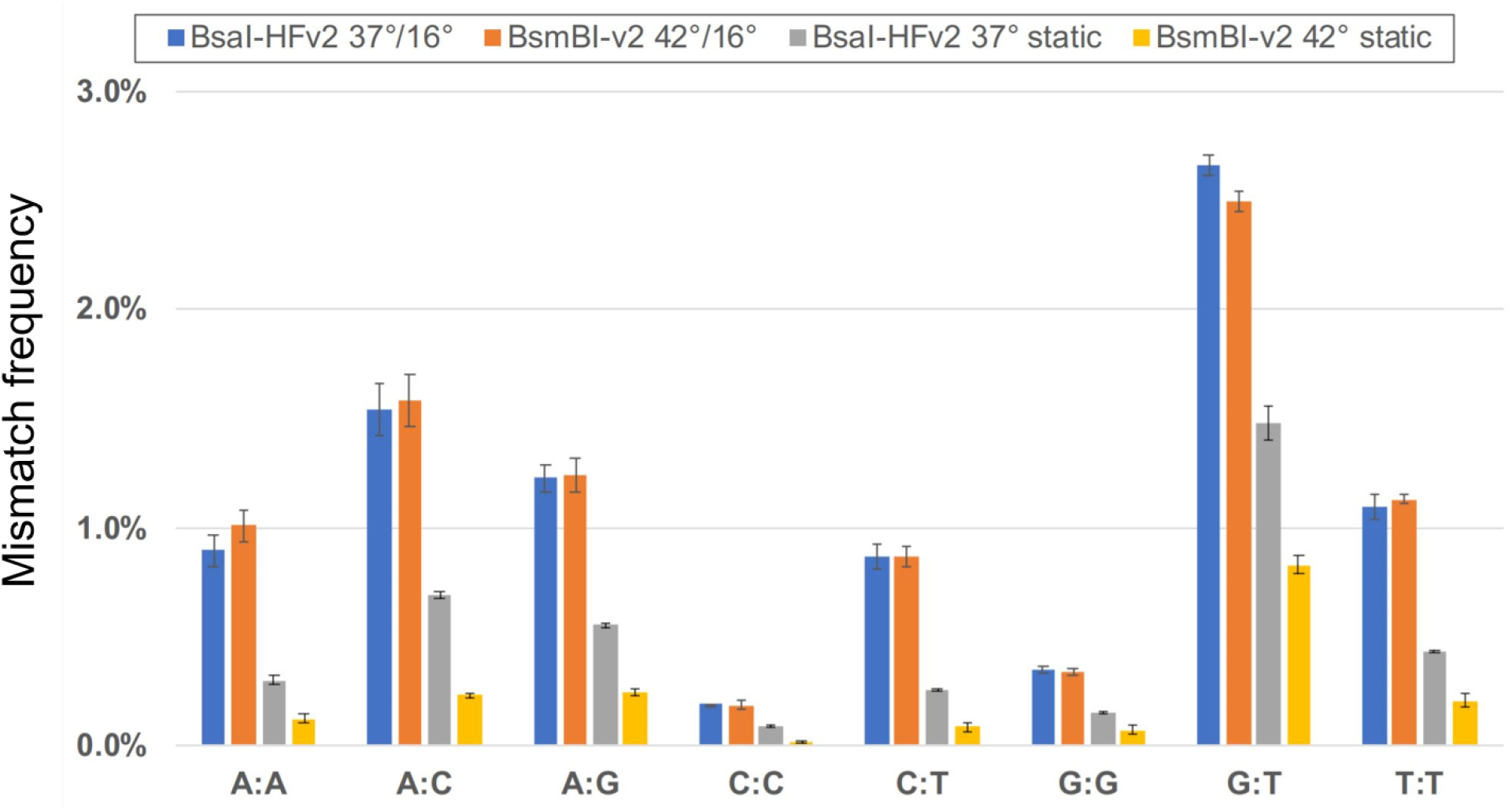
The impact of reaction temperature on nucleotide mismatch frequencies in assembly reactions with T4 DNA ligase and BsaI-HFv2 or BsmBI-v2. Mismatch frequencies for assembly reactions were grouped according to nucleotide mispair (A:A, A:C, A:G, C:C, C:T, G:G, G:T, T:T). Assembly reactions were carried out with T4 DNA ligase and either BsaI-HFv2 at 37°C (gray bars) or BsmBI-v2 at 42°C (yellow bars). For comparison, mismatch frequencies are shown for assembly reactions using traditional thermocycling protocols with T4 DNA ligase and either BsaI-HFv2 at 37°C and 16°C (blue bars) or BsmBI-v2 at 42°C and 16°C (orange bars); the mismatch frequencies for the thermocycling conditions has been reproduced here from our previous study [10]. The error bars depict the range between the maximum and minimum observed mismatch frequencies for two experimental replicates.

**Figure S2.**
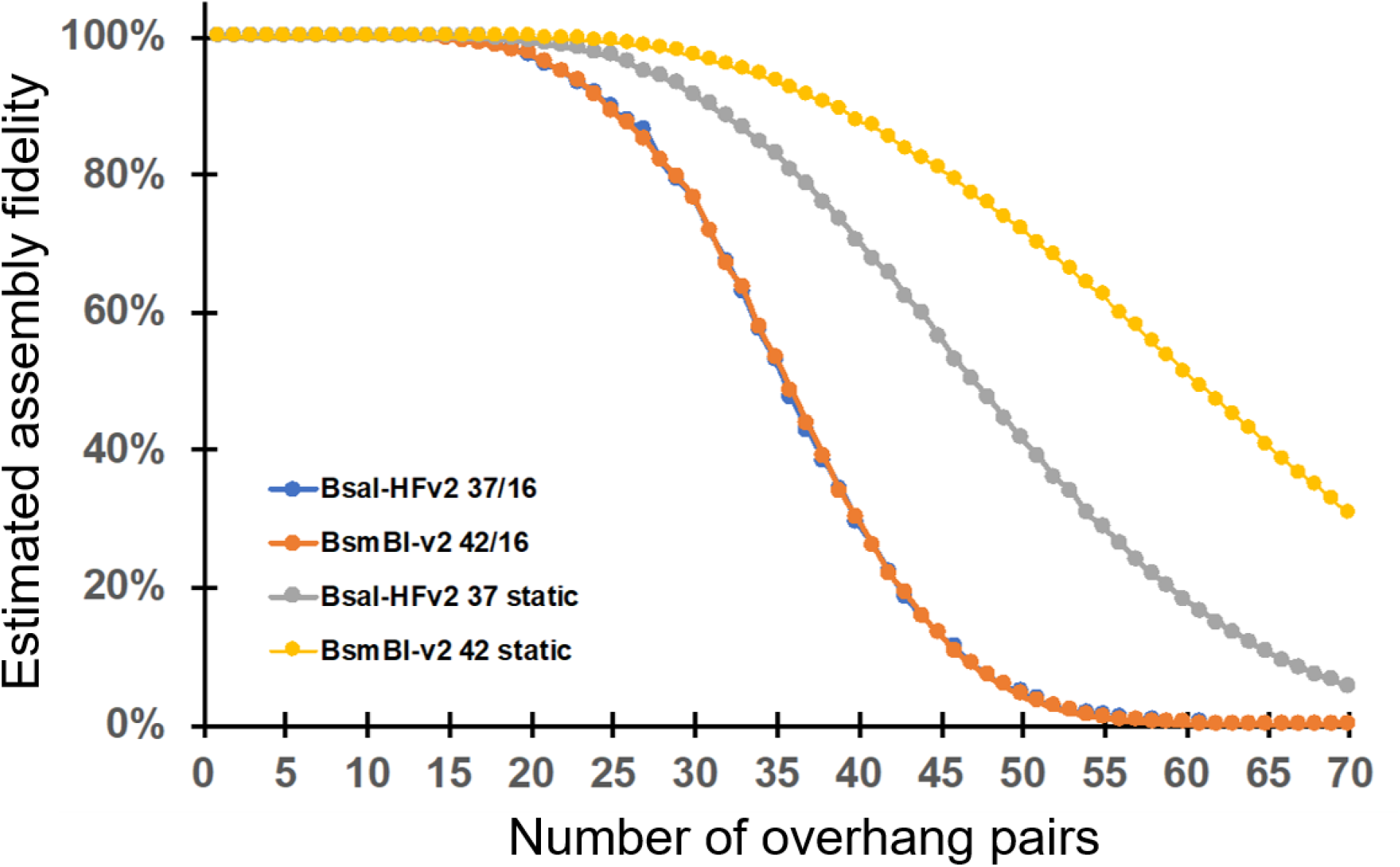
Golden Gate assembly fidelity predictions as a function of the overhang pairs in the assembly reaction. Fidelity estimations were carried out using the data from the multiplex DNA sequencing assay as described previously [10]. The estimated assembly fidelity of reactions containing up to 70 overhang pairs is shown for reactions using thermocycling conditions with T4 DNA ligase and BsaI-HFv2 (blue) or BsmBI-v2 (orange), or with T4 DNA ligase and BsaI-HFv2 at 37°C (gray) or BsmBI-v2 at 42°C (yellow).

**Figure S3.**
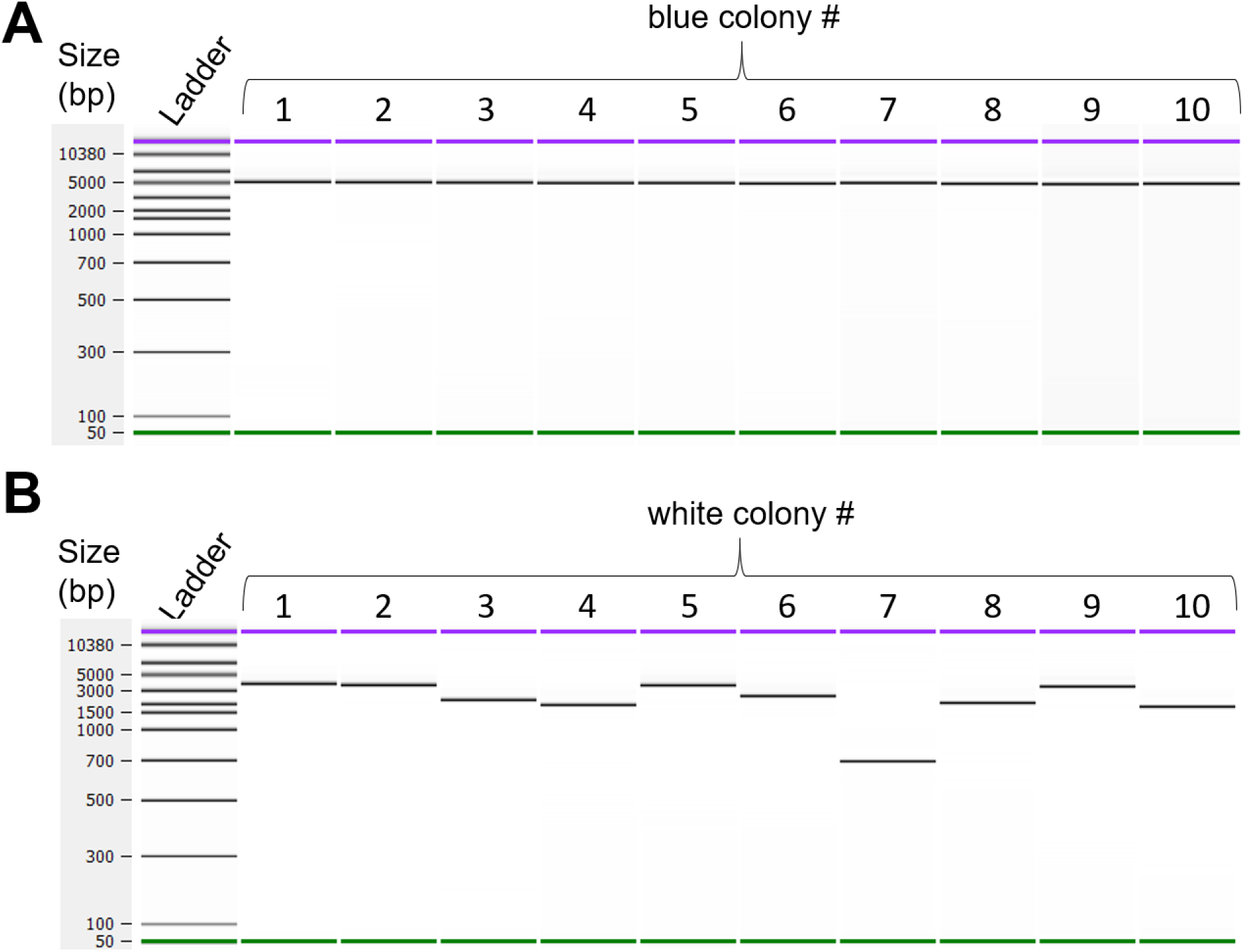
Verification of the 52 fragment *lac* operon cassette assembly. Plasmid DNA was isolated from colonies using the Monarch Plasmid Miniprep kit and subjected to PCR with amplification primers that flank the desired insertion site. As anticipated, (A) blue colonies contained inserts of the expected size for correct assembly of all 52 fragments, and (B) white colonies harbored constructs carrying truncated assembly products.

**Figure S4.**
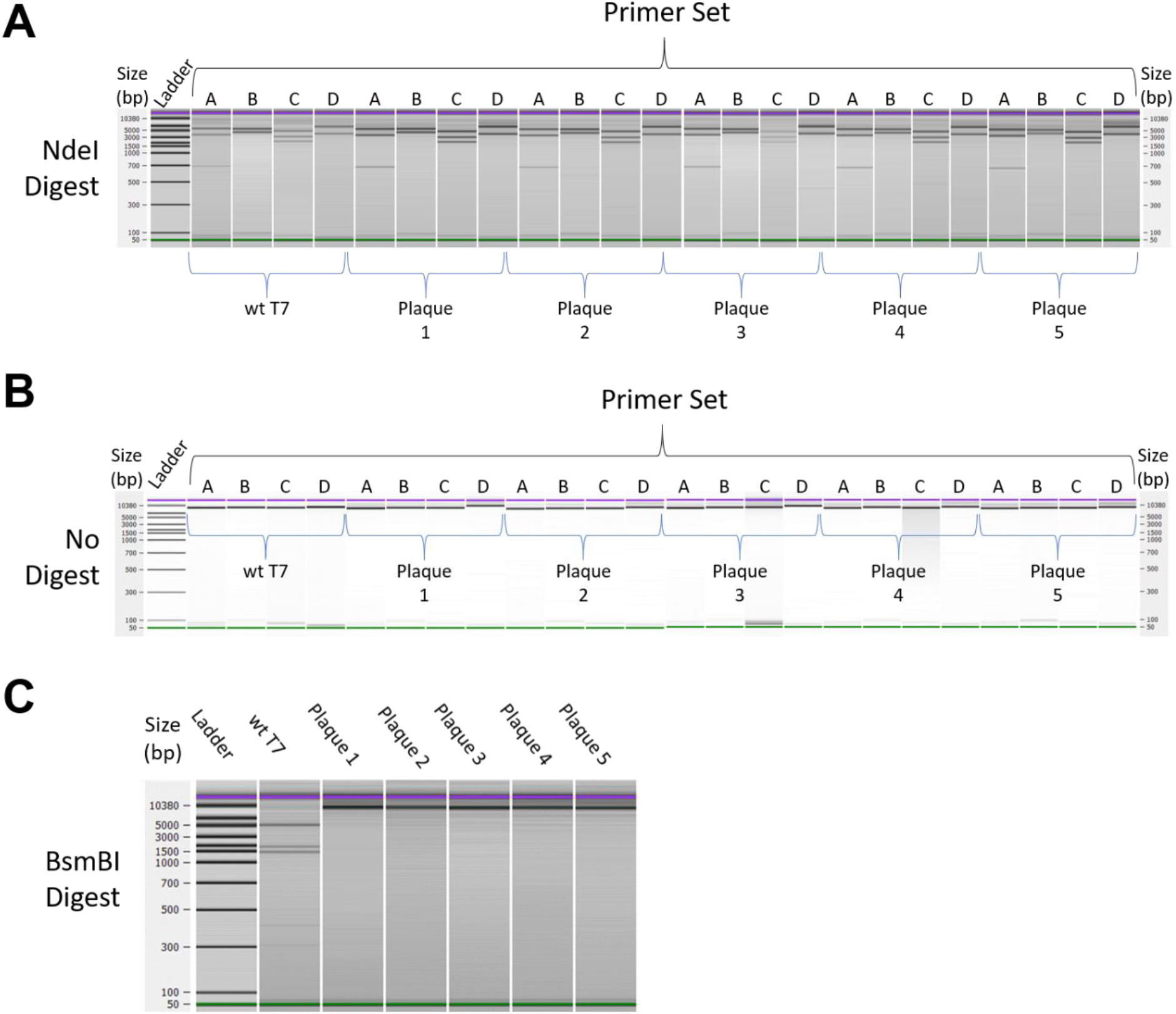
Verification of the 52 fragment T7 phage gDNA assembly. Plaque PCR was carried out using 4 sets of amplification primers (A-D) that together span the 40 kb phage genome. Amplicon lengths were resolved by Agilient Bioanalzyer 2100, using a DNA 12000 assay. Amplicons from 5 phage plaques were compared to the parental wt T7 phage genome after restriction enzyme digest with NdeI (A) or undigested (B). In all cases, the phage plaques produced a pattern identical to the parental wt T7 gDNA. (C) To confirm that the assembled genomes harbored the desired silent mutations to remove native BsmBI restriction sites, we also carried out amplicon digestion with BsmBI. We show that amplicon D from the parental T7 phage genome is digested by BsmBI, whereas amplicon D from the assembled phage genomes is inert to cleavage by BsmBI.

**Table S1. Ligation frequency for each overhang pair in assembly reactions with BsaI-HFv2 and T4 DNA ligase at 37°C**

**Table S2. Ligation frequency for each overhang pair in assembly reactions with BsmBI-v2 and T4 DNA ligase at 42°C**

**Table S3. 52 fragment *lac* operon cassette assembly fragments and overhang sequences Table S4. Primer sequences**

**Table S5. 52 fragment T7 phage assembly fragments and overhang sequences**

